# Reticulate leaf venation in *Pilea peperomioides* is a Voronoi diagram

**DOI:** 10.1101/2024.07.01.601217

**Authors:** Xingyu Zheng, Matthew Venezia, Elijah Blum, Ullas V. Pedmale, Dave Jackson, Przemyslaw Prusinkiewicz, Saket Navlakha

## Abstract

Reticulate leaf venation, characterized by the presence of loops, is a distinguishing feature of many flowering plants. However, our understanding of both the geometry and the morphogenesis of reticulate vein patterns is far from complete. We show that in the Chinese money plant (*Pilea peperomioides*), major veins form an approximate Voronoi diagram surrounding secretory pores known as hydathodes. We also propose a mechanistic model based on polar transport of the plant hormone auxin to produce Voronoi patterns. In contrast with classical models where veins directly connect auxin sources to sinks, our model generates veins that bisect the space between adjacent auxin sources, collectively forming loops. The paradigm change offered by this model may open the door to study reticulate vein formation in other species.

## Introduction

The quest to characterize natural forms in precise, mathematical terms has a history spanning centuries [1]. In general, these characterizations belong to three categories:

- <u>Partial descriptions</u>, which characterize some aspects of form, but are not comprehensive enough to reconstruct them. Examples include symmetry, allometric scaling (e.g., brain size scaling across mammals) [2, 3] and self-similarity (e.g., broccoli) [4].
- <u>Total descriptions</u>, which characterize an entire form. Examples include the spiral shapes of sea shells [5] and phyllotaxis (e.g., seed arrangement in the sunflower head) [6].
- <u>Generative models</u>, which explain how observed forms develop using local interactions. Examples include reaction-diffusion models of pattern formation (e.g., digit formation) [7] and Lindenmayer system models of branching structures (e.g., plant shoot architecture) [8].

The ultimate goal is to obtain a comprehensive characterization of form consistent with the partial descriptions, and a mechanistic model of its emergence. Here, we seek a comprehensive description and corresponding generative model for reticulated leaf veins.

Despite both their ubiquity in nature and accessibility to study, reticulate (i.e., closed loop) venation patterns have only been partially characterized, with few plausible generative models. Partial descriptions include the geometric relationships between branch radii and branch angles [9, 10]; the spatial distribution of veins across a leaf blade [11–13]; the hierarchical nesting and ordering of loops [14–16], including their sizes and shapes [17]; and how loops provide robustness to vein damage and flow fluctuations [18, 19]. Generative models of venation patterning at the molecular level are dominated by Sachs’ canalization hypothesis [20–22], which attributes the patterning of veins to positive feedback between the plant hormone auxin and its transporters. These models, however, typically produce tree-like branching patterns, as opposed to closed loops [23–27]. Together, these descriptions and models remain inadequate to explain the development of reticulated veins using plausible molecular mechanisms.

Voronoi diagrams [28–30] are geometric patterns that have been studied for centuries in both computational geometry and mathematical biology. A Voronoi diagram is a partitioning of space into polygons (i.e., closed loops), each surrounding a given center. This diagram has the property that all the points within a polygon are closer to that polygon’s center than any other center. For example, when partitioning a town (space) into school districts (polygons), a Voronoi partitioning guarantees that all students living within a district are closer to that district’s school (center) than any other school, thus minimizing distances traveled. While many biological structures found in nature also seem to resemble Voronoi diagrams [31], in most of these examples (e.g., giraffe skin patterns [32] or secondary veins of dragonflies [33]) only the polygon boundaries but not the centers are present (Supplementary Text Section 1). Thus, rigorously testing whether these examples are naturally occurring Voronoi diagrams is not straight-forward.

We discovered that in *Pilea peperomioides*, commonly known as the Chinese money plant, Voronoi diagrams are an approximate global descriptor of major vein reticulation. This is the first demonstration, to our knowledge, of the existence of Voronoi diagrams in plant venation systems, where both edges and centers are visible and functional. We also propose a generative model capable of approximating the observed Voronoi-like vein patterning using a plausible, auxin-flux patterning mechanism. This model offers an alternative to the canalization hypothesis and proposes a conceptually new direction for understanding the form and morphogenesis of reticulated networks in plants.

## Results

### *Pilea* as a model system for reticulate vein patterns

*Pilea peperomioides* (Fig. 1A) is an angiosperm from the *Urticaceae* family, native to southern China [34]. We studied the relationship between two structures found on *Pilea* leaves: hydathodes and major veins (Fig. 1B–C). Hydathodes are secretory pores and guttation sites found on the leaves of many plant species. These pores perform numerous functions, including releasing excess water to regulate hydraulic balance [35], removing harmful chemicals [35], and regulating immune responses against leaf-invading microbes [36]. Compared to stomata, hydathodes are roughly an order of magnitude larger, allow for uni-directional flow (as opposed to bi-directional gas exchange), and are static structures that remain open. In *Pilea*, the major veins, including both primary and secondary veins, form polygon-shaped closed loops surrounding hydathodes [37], whereas minor veins branch from major veins and connect directly to hydathodes (Fig. 1C–D; Supplementary Text Section 2).

**Fig. 1.**
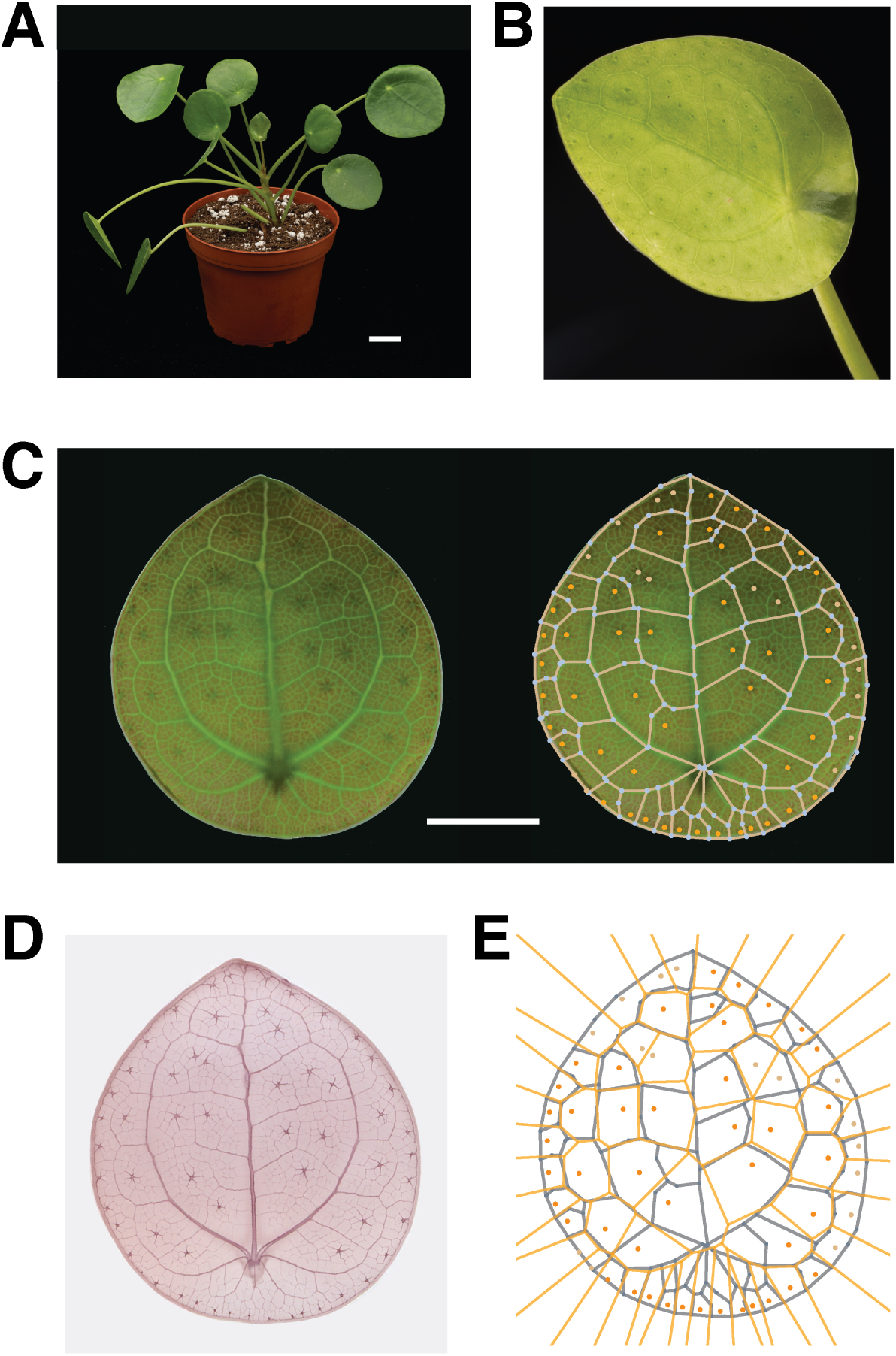
Major veins and hydathodes in *Pilea peperomioides* form a Voronoi-like diagram. (**A**) The *Pilea* plant (left); scale bar = 2 cm. (**B**) Close-up view of a young leaf (adaxial side up). (**C**) Example *Pilea* leaf under light microscopy (left); overlay of the leaf with the traced major vein network (white edges) and hydathodes (orange dots); scale bar = 2 mm. (**D**) Cleared scan of the same leaf, with vasculature and hydathode stained with safranin. (**E**) Overlay of actual major veins (blue lines) with an exact Voronoi diagram (orange lines) using hydathodes as the centers.

We imaged flattened *Pilea* leaves using light microscopy and developed a semi-automated computational pipeline to extract the two-dimensional positions of all hydathodes, as well as the network structure formed by major veins (Fig. 1E and Fig. S1; Materials and Methods). We analyzed 34 leaves from 6 *Pilea* plants, grown in indoor conditions (Materials and Methods). Most minimal polygons (i.e., polygons that contains no other polygons) of major veins contained exactly one hydathode (73.11%), and the remaining polygons contained either zero (10.26%) or *>* 1 hydathodes (16.63%), which we discuss later. Polygons in each leaf had a wide spread of sizes and aspect ratios (Table S1).

### A geometric framework to quantify Voronoi patterning

Given a set of *n* centers in two-dimensional space, a Voronoi diagram consists of a set of *n* polygons 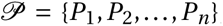, each enclosing a single center, such that all points *p* within polygon *P_i_* are closer to center *i* than any other center *j ̸= i* :

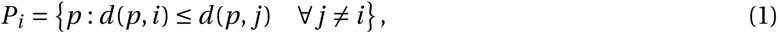

where *d* (*·*, *·*) denotes Euclidean distance. The boundaries of polygons are formed along points that are equidistant to adjacent centers.

Due to stochasticity and developmental constraints, it is unlikely that a naturally formed venation patterning will perfectly satisfy the Voronoi property in Equation (1). Thus, we developed three tests to quantify how close a given patterning is to matching the Voronoi property. These tests, described next, measure: I) how well local geometric relationships between hydathodes and veins follow those of an ideal Voronoi diagram; II) how well vein locations can be predicted given only hydathode locations as input; and III) how well hydathode locations can be predicted given only vein locations as input. We also applied these tests to tessellations that would be generated if each center (i.e., the hydathode) were located at three alternative reference locations within a polygon: 1) the centroid of the polygon (i.e., its gravitational center), which is often used as a proxy for center locations when centers are absent [38, 39]; 2) the mid-point of the polygon: produced by sliding a horizontal line though the polygon in the vertical direction, stopping when the line is mid-way through the polygon, and finding the mid-point of the line segment bounded by the polygon edges; and 3) random points picked within the polygon.

### *Pilea* hydathodes and major veins form a near-exact Voronoi diagram

#### Voronoi Test I: Angle and distance conditions

A corollary of the Voronoi property (Equation (1)) is that the line segment connecting the centers of adjacent polygons should perpendicularly bisect the shared edge of the two polygons (Fig. 2A). This means that the angle (*θ*) formed by the line segment and the shared boundary should be 90 degrees, and that the two distances (*d*_1_, *d*_2_) from each adjacent center to the shared boundary should be equal. We defined the angle error to be Δ*θ = |*90*−θ|*, and the (normalized) distance error to be Δ*d = |d*_1_ *− d*_2_*|*/(*d*_1_ *+ d*_2_). Smaller errors indicate a closer match to an ideal Voronoi diagram. We computed the angle and distance errors for all 1836 adjacent pairs of major vein polygons across all 34 leaves (Table S1) that each contained a single hydathode.

**Fig. 2.**
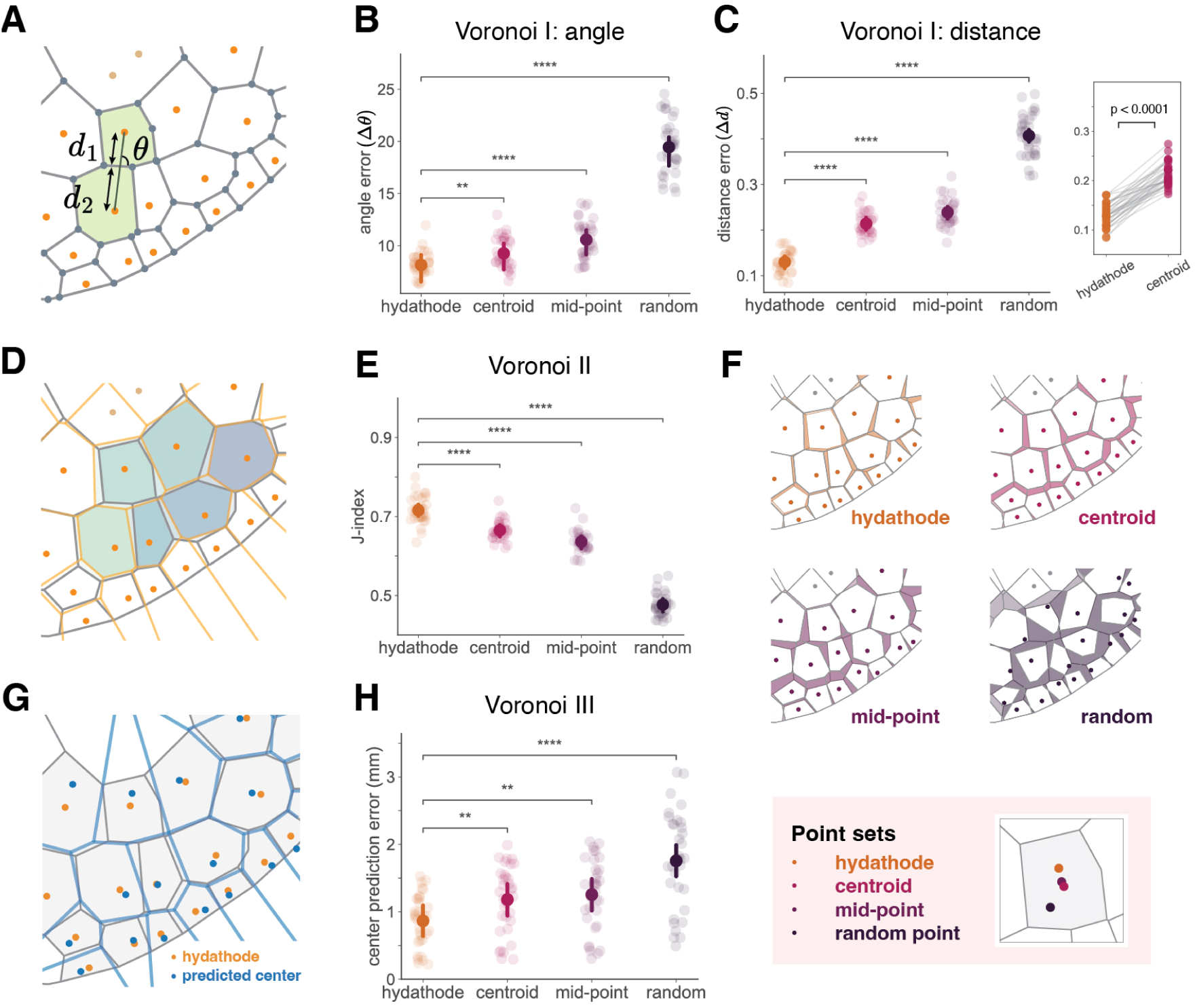
The Voronoi diagram approximation in *Pilea* is supported by three statistical tests. (**A**) Schematic for the Voronoi I test, applied to each neighboring polygon pair. (**B**) Voronoi I test result (*n =* 34) for the angle error (Δ*θ*). The mean and 95% interval of the estimated mean of the posterior is plotted in the foreground; see Materials and Methods for significance calculation. (**C**) Voronoi I test result (*n =* 34) for distance error (Δ*d*). Inlet highlights pairwise comparison between errors computed with hydathodes vs. centroids. (**D**) Schematic for Voronoi II test. Overlay of the *Pilea* major vein graph (grey) and Voronoi diagram edges (orange) using hydathodes as the centers. Unions and intersections of corresponding polygons in the two tessellations are shaded and used to calculate the Jaccard Index. (**E**) Voronoi II test results (*n =* 34). (**F**) Visualization of the pattern mismatch between *Pilea*’s major vein polygons and the Voronoi diagrams generated by hydathodes vs. other reference point sets. (**G**) Schematic for Voronoi III test. Overlay of the *Pilea* major vein graph (grey), hydathodes (orange) with the predicted center location (blue dots), and the Voronoi diagram (blue edges) generated using the predicted centers. (**H**) Voronoi III test results (*n =* 33).

The errors induced when using the hydathode locations as Voronoi centers were less than when using the three other reference locations (Fig. 2B–C). Specifically, the angle error for hydathodes was 8.23*±*1.13 degrees (mean *±* standard deviation), significantly smaller than 9.35 *±* 1.47 for centroids, 10.65 *±* 1.82 for mid-points, and 19.51*±*2.48 for random points (Fig. 2B) (Materials and Methods; Table S2). Similarly, the distance error for hydathodes was 0.13*±*0.02, which was significantly less than that of the other reference locations: 0.21 *±* 0.02 for centroids, 0.24 *±* 0.03 for mid-points, and 0.41 *±* 0.04 for random points (Fig. 2C; Table S2). Thus, the spatial arrangement of hydathodes and major vein polygons closely followed the local geometry of a Voronoi diagram.

#### Voronoi Test II: Predicting veins given hydathode locations

If a true Voronoi diagram were constructed given actual hydathode locations as input, how much overlap would there be between the true Voronoi polygons and the actual polygons formed by major veins? To quantify this, we measured the average area overlap between the true Voronoi polygons and corresponding observed (major vein) polygons (Fig. 2D):

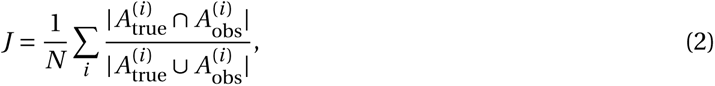

where *N* is the number of single-hydathode polygons on the leaf, and 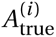 and 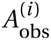 are sets of points in the polygon for center *i* in the true Voronoi diagram and the corresponding major vein polygon, respectively. This measure, called the Jaccard index, ranges from 0 to 1, with larger values indicating more overlap.

The overlap between true and observed polygons was better when using hydathodes as centers, compared to when using the other reference locations (Fig. 2E). Specifically, the overlap for hydathodes was 0.72 *±* 0.03, which is significantly higher compared to 0.67 *±* 0.02, 0.64 *±* 0.03, and 0.47 *±* 0.03 for centroid, mid-point, and random, respectively (Table S2). These differences are visually pronounced (Fig. 2F), especially at the periphery of the leaf blade, where the true and hydathode-predicted boundaries are particularly well-aligned, whereas the corresponding polygons from the reference points are not.

To measure how much biological noise would be required to match the observed deviation from a Voronoi diagram, we added random noise at each true Voronoi vertex (i.e., at locations where two polygon boundaries intersect) and calculated the Jaccard index between the true (noise-free) Voronoi polygons and those formed after noise was added. The overlap between the noise-free and noisy Voronoi diagrams matched the observed overlap (0.72) when about 15% noise was added (Fig. S2A, B). Thus, the observed deviations were commensurate with the ideal Voronoi with a small amount of noise.

To test if an alternative spatial tessellation model can generate polygons similar to *Pilea* major vein polygons, we applied a variant of the *k*-*d* tree (Materials and Methods), which recursively generates rectangular cells surrounding hydathodes until all pairs of adjacent hydathodes are bisected by a cell edge (Fig. S2C–E). This model uses a hierarchical spatial sub-division principle shared with the venation process [40–42], and it generates perpendicular bisections between adjacent cell edges and centers. However, the overlap between the *k*-*d* tree tessellation and the major vein polygons was 0.40 *±* 0.03, which was on par with the overlap for random points (0.47). Thus, Voronoi diagrams are a more likely candidate model than a *k*-*d* tree to describe the relationship between hydathodes and major veins.

#### Voronoi Test III: Predicting hydathode locations given veins

Given the polygons formed by major veins, how well can we predict the locations of the enclosed hydathodes? This is called the “inverse Voronoi” problem [43], and our goal here was to solve for the center locations that minimize the angle and distance errors for a given tessellation (Materials and Methods) [44].

We found that the optimal Voronoi center locations were significantly closer to the actual hydathode locations than any of the three reference locations (Fig. 2G–H). Specifically, hydathode locations were 0.87 *±* 0.38 mm from the true Voronoi center locations, compared to 1.18 *±* 0.50 mm for centroid, 1.25 *±* 0.53 mm for mid-point, and 1.76 *±* 0.72 mm for random points (Table S2). This suggests that hydathodes are located near the true Voronoi centers of major vein polygons.

In general, these three Voronoi tests can be applied to determine how close a natural Voronoi-like diagram is to a true Voronoi diagram. To verify their stringency, we applied them to another biological spatial tessellation: the air chambers (polygons) surrounding air pores (centers) on leaf-like structures (thalluses) in the liverwort *Marchantia polymorpha* (Fig. S3). We found that air pore locations deviated more from a true Voronoi diagram compared to centroid locations, and the Voronoi test errors for the liverwort thallus were larger than those in *Pilea* (Table S2, S3). This suggests that *Marchantia* air chambers and pores do not form a Voronoi diagram, and that our tests indeed have the statistical power to distinguish Voronoi-like patterns from other non-Voronoi spatial tessellations (Supplementary Text Section 6).

### The *Pilea* Voronoi property is robust across stress conditions

We next tested how robust the Voronoi property was to changes in growth caused by exposing *Pilea* plants to light or heat stress (Fig. 3A), conditions which impact hydathode morphology [35, 45] and vein patterning [46], respectively. We grew 6 plants in each of three stress conditions, namely shade, high-light intensity, and high-temperature, and collected 128 new leaves that emerged during a five-week treatment period (Materials and Methods). *Pilea* exhibited various phenotypic changes in response to these conditions, especially changes in leaf morphology (Fig. S4A-E; Supplementary Text Section 3). For example, hydathode sizes varied significantly across conditions (Fig. 3B): in high-temperature, hydathodes were significantly larger than in controls, suggesting that hydathode size is regulated to manage water loss. These conditions also led to variation in leaf shape, color, and size, though not in the number of hydathodes (Table S1; Supplementary Text Section 3). The histogram of hydathode counts in each condition followed a right-skewed unimodal distribution with means that were not significantly different (Fig. S4F–G). Moreover, the number of hydathodes per leaf could be described by a simple Poisson distribution centered at the average of all samples combined (Fig. 3D). While there was stochasticity in exact hydathode locations across leaves in different conditions, the underlying spatial distribution of hydathodes was similar (Fig. 3C; Supplementary Text Section 4).

**Fig. 3.**
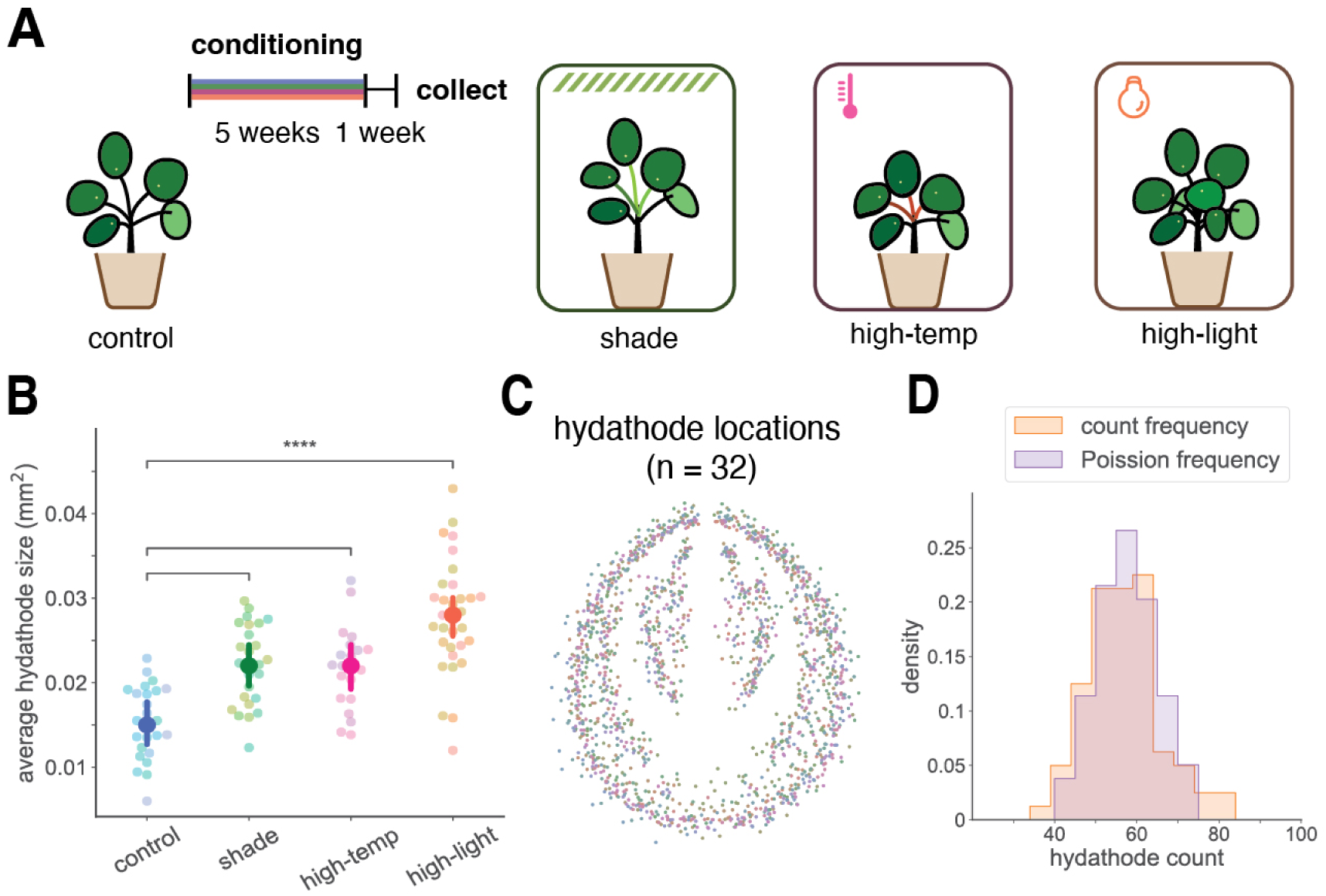
The *Pilea* Voronoi property is preserved under environmental stress. (**A**) Schematic of environmental conditioning experiments. *Pilea* plants were exposed to four conditions: control (indoors with indirect natural light), shade, high temperature, and high light intensity. (**B**) Mean hydathode size in control and treatment conditions (*n =* 20 each). The mean and 95% interval of the estimated posterior for the mean hydathode size is plotted in the foreground, and the colors of the scatter plot in the background are coded by plants. (**C**) Overlay of hydathode locations from 32 samples over all treatment conditions. (**D**) Relative frequencies of the hydathode count distribution (*n =* 80) and of the Poisson distribution with the same mean.

Despite these changes in leaf morphology, hydathodes and major vein polygons still formed near-exact Voronoi diagrams (Fig. S5) according to the three Voronoi tests. Moreover, the level of deviation from being a true Voronoi diagram was low for the stressed plants, similar to the controls (Table S2, S3). The preservation of Voronoi-like patterning in the presence of stress-induced variation in the number and location of hydathodes implies that this pattern is not pre-specified in development, and that a local, self-regulatory mechanism might be at play.

### Voronoi-like vasculature patterns in *Pilea* leaves can be patterned by auxin

A key remaining question is the molecular mechanism giving rise to the observed relation between hydathodes and vascular patterning in *Pilea*. The prevalent theory for vein formation is canalization, proposed by Sachs more than half a century ago, which attributes the patterning of vascular strands to a positive feedback between auxin flow and the polarization of auxin transporters [20–22]. Predating molecular evidence, Mitchison [23, 47, 48] used a computational model to demonstrate that, if this feedback is strong enough, auxin flowing from auxin sources to sinks will become channeled into gradually narrowing paths with high auxin flux (canals), the precursors of veins. The subsequent discovery of auxin efflux carriers (PINs), and the visualization of PIN and auxin response reporter proteins, lent further credence to the hypothesis that auxin and its transporters are central to vein patterning [49–52]. Follow-up computational models further confirmed that strong feedback between auxin flux and PINs can reproduce midvein formation [24] and open branching patterns [25]. However, the Voronoi-like relation between hydathodes and major vascular strands in *Pilea* leaves is at odds with canalization. Hydathodes are sites of high auxin concentration [53–55] and thus likely act as auxin sources, yet major veins run between the hydathodes, separating rather than connecting them. This raises the question of whether an alternative mode of interaction between auxin and PINs could explain the patterning of major veins in *Pilea*.

In the absence of sufficient molecular-level data, we addressed this question using computational models. We hypothesized that the patterning of main veins in *Pilea* is driven by auxin, as in other plants, but the altered parameters of this process make it distinct from canalization. Our model was inspired by Feugier et al. [25], who analyzed different forms of feedback between auxin flow and the polar allocation of its transporters. They showed that, if the allocation of auxin carriers in response to the auxin flux is stronger than linear (e.g., quadratic, as postulated by Mitchison [23]), the resulting feedback will lead to the formation of canals transporting auxin from sources to sinks. In contrast, if the feedback is weaker — linear or decreasing — auxin will flow away from its sources without forming canals. Weak polarization was proposed as a mechanism that positioned primordia in a model integrating phyllotaxis and vascular patterning in the shoot apical meristems of *Arabidopsis* [56], and as a mechanism that directed vascular strands (subsequently patterned using strong canalization) towards sinks in the meristems of *Brachypodium* [26]. Here we show that weak polarization can pattern vascular strands by itself as well. These strands do not connect auxin sources to sinks, but divide the space between the sources similarly to the major veins partitioning the space between hydathodes in *Pilea*. We assumed the polarization model with directional fluxes [57] due to its biochemical plausibility [58], and we implemented it in the version ignoring intercellular spaces [26] for simplicity.

The model simulates the dynamics of auxin distribution in grids of square cells. For theoretical investigations we considered 1D strands of cells and 2D rectangular grids; for the models of leaves, we clipped a 2D grid to the shape of a leaf and assumed positions of hydathodes obtained from actual leaves. Hydathodes act as the sources of auxin and, once activated, maintain constant, high auxin concentration. The concentration *c_A_* of auxin in an arbitrary ground (non-hydathode) cell *A* changes according to the equation:

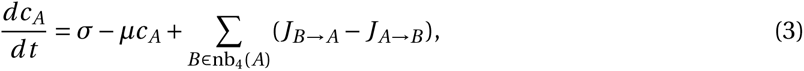

where *σ* is the background rate of local synthesis of auxin, *µ* is the rate of auxin turnover, and *J_B→A_*, *J_A→B_* represent unidirectional auxin fluxes due to its polar transport to and from cell A. These fluxes are summed over all cells *B* that share an edge with *A*, i.e., that lie in the 4-neighborhood nb_4_(*A*) of *A*. The unidirectional flux *J_A→B_* between any pair of neighboring cells *A*, *B* is calculated as:

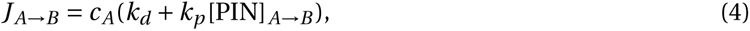

where *k_d_* is the rate of auxin diffusion through the wall between cells *A* and *B* (including a possible role of plasmodesmata [59]), and *k_p_* is the rate of polar transport from *A* to *B* by PIN proteins located on the membrane of cell *A* abutting cell *B*. The abundance of this protein, [PIN]*_A→B_*, is the net effect of PIN cycling from the cell interior to the membrane (exocytosis) and from the membrane back to the interior (endocytosis):

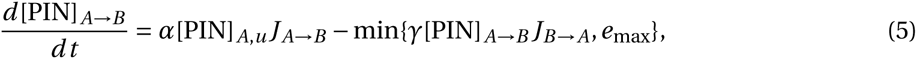

where [PIN]*_A_*_,*u*_ is the abundance of currently unallocated PIN in the interior of cell *A*, *α* is the auxin outflux based exocytosis rate of PIN, *γ* is the auxin in-flux based endocytosis rate, and *e*_max_ is the maximum endocytosis rate.

Finally, we assume that [PIN]*_A_*_,total_, the total abundance of PIN in each cell *A*, is constant, thus

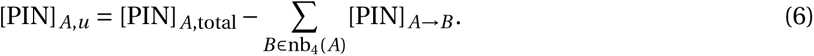

The operation of this model can be illustrated by a sequence of simulations, which gradually activate different model components (Fig. 4); parameter values used in the simulations are listed in Table S4. We first consider a one-dimensional row of cells, in which the leftmost cell has just been activated as an auxin-producing hydathode and will maintain high constant auxin concentration throughout the simulation; the remaining ground cells have low initial concentration. With a purely diffusive transport (*k_d_ >* 0, *k_p_ =* 0 in Equation (4)), auxin concentration gradually increases in all ground cells, tending to the concentration of the hydathode. This process begins with the hydathode and progresses away from it (Fig. 4A). At no stage does a concentration peak emerge that could potentially trigger a localized differentiation of vascular cells. This dynamic is drastically changed in the presence of polar auxin transport (*k_p_ >* 0) with feedback between auxin flow and the allocation of auxin carriers. For a wide range of parameter values *γ ≫ α >* 0 (Equation (5)), a wave of high auxin concentration emerges, propagating away from the hydathode at an approximately constant speed (Fig. 4B). With two hydathodes arising simultaneously, the generated waves collide, producing a maximum of auxin concentration half-way between the hydathodes (Fig. S6A, highlighted by yellow box). The auxin concentration in this maximum quickly rises, which reverts polarization of the adjacent cells making the maximum disappear (Fig. S6), but it can be stabilized by various mechanisms. In the model we have imposed a cap, *c*_max_, on auxin concentration (Fig. 4C); a more complex mechanism, for example involving the drainage of auxin through the petiole, may plausibly operate in actual *Pilea* leaves.

**Fig. 4.**
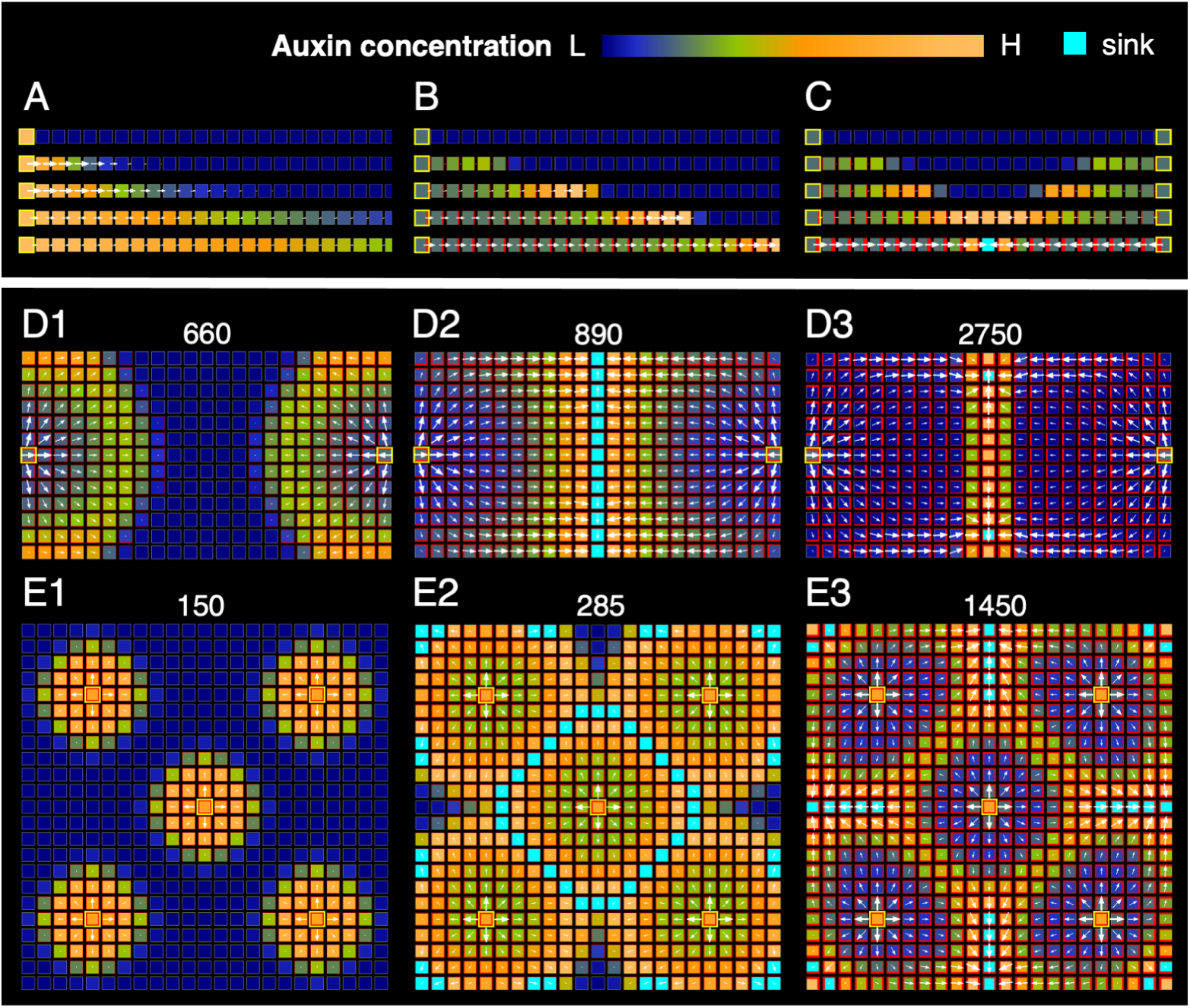
Principle of Voronoi diagram formation by the auxin-driven patterning with unilateral fluxes. Auxin sources are highlighted by yellow boxes. Auxin concentration is indicated by the cell color. The color scheme used is indicated by a color map (top right) ranging from dark blue (low auxin, L) to pale orange (high auxin, H). Cells (sinks) that reach the maximum concentration cap, *c*_max_, are colored in cyan. PIN concentration at the cell membranes is indicated by the width of red lines along the cell edges, and auxin fluxes are represented by the direction and size of white arrows. See Table S4 for model parameter details. (**A**) Snapshots of a simulation of pure auxin diffusion in a semi-infinite row of cells. Auxin gradually infiltrates consecutive cells without forming new maxima. (**B**) Same as (A), with polar auxin transport. An auxin maximum emerges, forming a wave that propagates away from the source. (**C**) Same as (B), in a finite-length row of cells with auxin sources at both ends. Colliding waves generate a concentration peak, which is stabilized by a concentration cap (cyan). (**D**) Same as (C), simulated on a rectangular grid with no-flux boundary conditions, and an additional cap on the rate of endocytosis. Auxin concentration waves propagate (D1) and collide half-way between the cells (D2), forming a ridge that persist for some time after collision (D3). The numbers indicate frame numbers generated by the simulation. (**E**) Same as (D), on a square grid with 5 auxin sources. Waves of high auxin concentration collide along ridges that bisect the space between the sources, producing a Voronoi diagram.

The generation of auxin maxima partially extends to two dimensions. In this case, wave fronts are approximately circular and collide along a linear ridge (Fig. S7 and Video S1). This ridge is unstable, and even with a cap on auxin concentration, PIN quickly repolarizes away from the ridge center, redirecting auxin flow toward emerging discrete maxima (Fig. S8 and Video S2). A properly chosen cap on the rate of endocytosis (parameter *e*_max_ in Equation (5)) significantly slows down the repolarization while preserving the preceding wave propagation and collision phases (Fig. 4D, Fig. S9 and Video S3). A more complex mechanism, for example involving the CUP-SHAPED COTYLEDON2 (CUC2) protein known to control PIN reorientation in *Arabidopsis* leaves [60], may operate in actual *Pilea* leaves. In the presence of several hydathodes, the ridges form loops that surround the individual hydathodes and, with the hydathodes arising approximately at the same time, bisect the space between them (Fig. 4E and Video S4). The patterning process is thus an auxin-based implementation of the fundamental process generating Voronoi diagrams by colliding waves [30, 61], in which hydathodes act as the polygon centers.

To examine the model’s capability to simulate actual venation patterns, we digitized the contours and hydathode positions of sample *Pilea* leaves, and simulated the patterning process with these input data (Fig. 5A and Video S5; Fig. S10 and Videos S6–S9). Overlaying the resulting pattern with that of a true Voronoi diagram generated using the same hydathode positions confirmed that the auxin-driven model produced a very close approximation of a Voronoi diagram (Fig. 5B). Superimposing the generated pattern on a cleared leaf image thus reflected both the similarities and differences between exact Voronoi diagrams and the actual major vein patterns (Fig. 5C). To characterize them further, we divided the real leaf (Fig. 5D) and the model pattern (Fig. 5E) into three zones: peripheral (a, unshaded), intermediate (b) and central (c–e). These zones were conspicuous in both the cleared leaf and the model, and had similar shapes. In the peripheral and intermediate zones (a, b), where the hydathodes were distributed relatively regularly along the contour of the leaf edge or a primary vein arc, the polygons of the model and cleared leaves coincided closely. In the central regions (c, d) of the actual leaves, some polygons included several hydathodes not separated by major veins, whereas simulated patterns adhered strictly to the Voronoi diagram. In spite of these differences, the capability of weak polarization to generate vascular patterns with loops was remarkable.

**Fig. 5.**
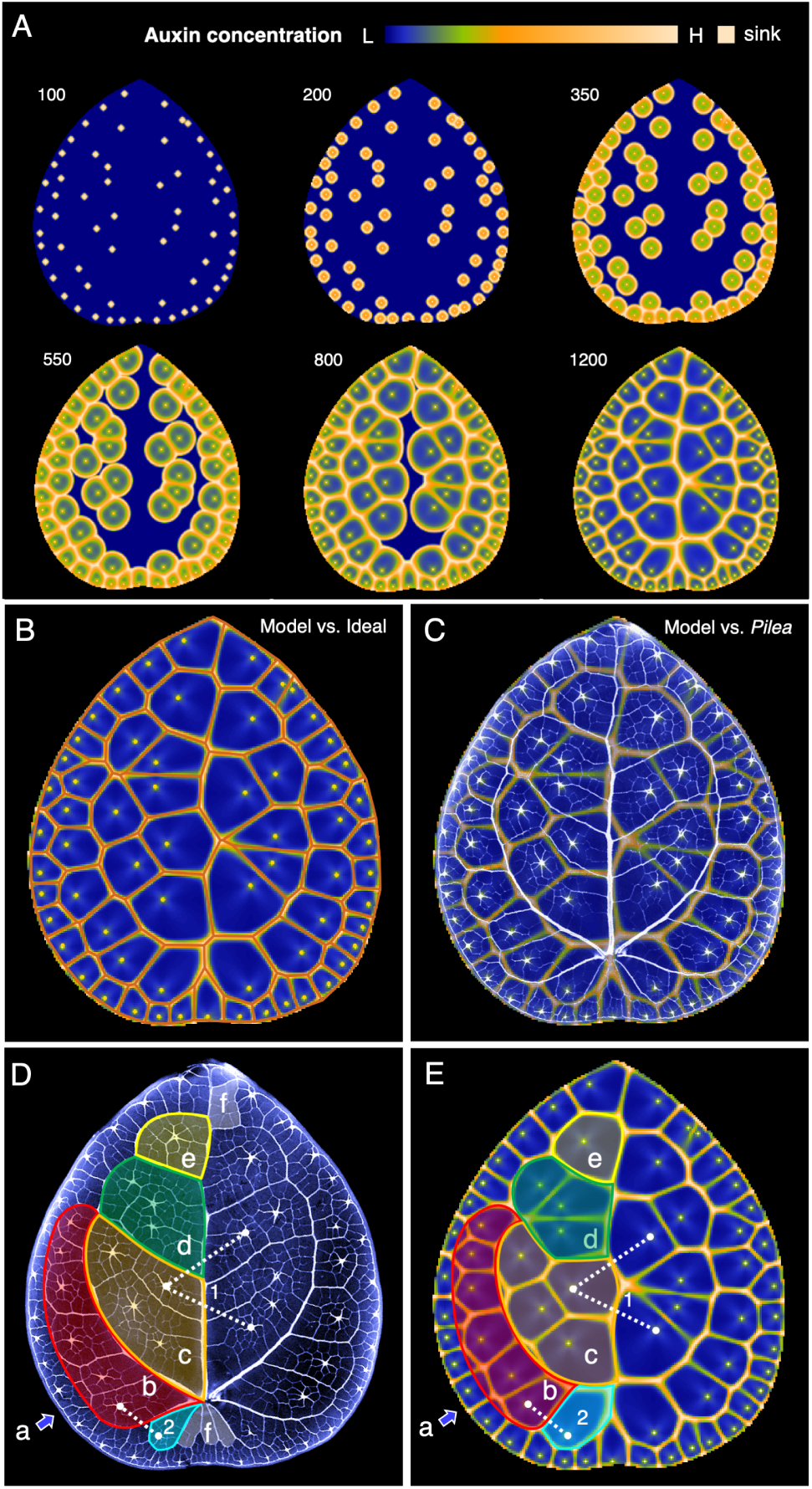
Simulated auxin-driven patterning of *Pilea* leaves. Simulation was performed on a 200 *×* 220 grid of cells, clipped to the contour of an actual leaf. Hydathode positions were obtained by digitizing the leaf. (**A**) Snapshots of the simulation illustrating the patterning process. Wavefronts originating at the hydathodes propagate and collide, forming high-concentration ridges as in Fig. 4E. The color map indicates auxin concentration from low (L) to high (H); the same color was used to represent *c*_max_ and the highest auxin concentration. The numbers indicate frame numbers generated by the simulation. See Table S4 for model parameter details. (**B**) Overlay of the model-generated pattern (yellow) with an ideal Voronoi diagram (red) computed using hydathodes as the polygons centers. (**C**) Overlay of the pattern generated by the auxin-driven model and the cleared *Pilea* leaf. **(D, E)** Cleared and modeled leaves annotated to facilitate comparisons. a: zone on the leaf periphery; b: zone sandwiched between zone a and the primary lateral vein; c–e: three conspicuous loops adjacent to the midvein; f: loops without hydathodes in the cleared leaf. Additional annotations highlight differences in vein orientation between the modeled and cleared leaves (dotted lines (1 & 2)), and differences in polygon shapes and sizes (cyan).

## Discussion

Reticulate leaf venation patterns, defined by the presence of loops, are ubiquitous in flowering plants, yet their geometry has so far eluded concise characterization. We have shown that the patterns of major veins in *Pilea* leaves approximate Voronoi diagrams, with hydathodes acting as the Voronoi polygon centers. Given the diverse and complex reticulate vascular patterns observed in nature, it was unexpected to find an example where the pattern can be explained by such a simple geometric principle.

We have also proposed and modeled a biologically plausible mechanism approximating the observed patterns. Similar to canalization, which has dominated the understanding of vascular patterning in plants for the last 50 years, our model attributes the patterning of veins in *Pilea* leaves to auxin. Canalization, however, creates channels of high auxin flux that connect auxin sources to sinks, and these channels primarily form open, tree-like vascular patterns. In contrast, our model generates auxin concentration waves that propagate away from hydathodes and form crests upon colliding. These crests pattern vascular strands that assemble into closed loops surrounding each hydathode, thus tessellating the leaf surface into Voronoi polygons.

Our model leads to the following predictions, which can potentially be tested by developing experimental methods in *Pilea* to analyze the dynamics of auxin propagation and the polarity of PIN homologs in developing leaves.

1. Hydathodes are likely sites of auxin biosynthesis [53–55], and thus disruption of auxin synthesis (e.g., via ablating or mutating hydathode structures [62]) at these sites would disrupt the Voronoi pattern of *Pilea* major veins.
2. Auxin concentration maxima, coinciding with the hydathode locations, are patterned before the surrounding major veins.
3. During the establishment of major veins, the efflux auxin transporter protein of *Pilea* (i.e., the *Pilea* version of PIN) are polarized away from hydathodes (Fig. 4E), creating waves of auxin concentration that collide, forming crests.

While the degree to which *Pilea* vasculature approximates a Voronoi diagram is striking, there remain patterning aspects not accounted for in our model. Future models that include these aspects may further reduce the following discrepancies between model-generated vein patterns and the *Pilea* realization.

1. In our model, the midvein zig-zags due to the asymmetry of hydathode distribution between the left and right sides of the blade, and it is patterned last because of the relatively large distances between hydathodes in the central zones on both sides of the leaf. In contrast, in *Pilea* leaves, the midvein is relatively straight and differentiates first, prior to hydathode formation (Table S5– Leaves). This discrepancy suggests that the midvein is patterned by a distinct process during leaf initiation, as is the case in *Arabidopsis* [51] and tomato [63] leaves.
2. In regions near the midvein and the attachment point to the petiole, there are discrepancies in the shapes and sizes of model-generated polygons vs. the vascular polygons in *Pilea*(Fig. 5C and Fig. S11; Supplementary Text Section 5), as well as deviation from the one-center one-polygon rule; most *Pilea* polygons with zero or multiple hydathodes lie in these regions. These discrepancies may be attributed to the fact that our model ignores the role of the petiole, which likely acts as a global sink for auxin in the leaf.
3. Our model only accounted for the patterning of major veins, whereas *Pilea* vascular polygons also include minor veins, which connect the hydathodes to the major veins. Minor veins may form by canalization, with major veins acting as local auxin sinks. Patterning processes combining weak and strong polarization have been proposed [26, 56], so it is possible that they are also integrated during the patterning of *Pilea* leaves.
4. Our model assumed the leaf size and shape were fixed and that hydathodes (as auxin sources) emerged synchronously from pre-specified locations. In reality, differentiation and maturation of leaf tissues, including that of hydathodes, follow a basipetal sequence in early development (Table S5– Leaves, Hydathodes). Future models that incorporate leaf growth and hydathode emergence may shed light on the degree to which this developmental dynamics affects the process of Voronoi pattern formation.

Our model may extend to reticulate patterning in other species with laminar hydathodes, where major veins surround hydathodes and connect to them via minor veins. Examples include *Ficus formosana* [64], *Ficus diversifolia* [65], and many other species in *Urticaceae*, the nettle family, to which *Pilea* belongs [66]. While the reticulate venation patterns found in these species may adhere to Voronoi geometry less strictly than in *Pilea*, the wave collision mechanism suggested by our model may potentially be adapted to generate these patterns by varying parameters that affect dynamics, such as the timing and quantity of auxin production in the hydathodes. For species with reticulate veins that do not have laminar hydathodes, it is nevertheless possible that auxin is synthesized at discrete sites distributed over the leaf blade, serving an analogous role to the hydathodes [67]. The combination of distributed local auxin maxima and weak polarization may be a general theme that gives rise to reticulation, forming an alternative to, or complementing, previous hypotheses on reticulate pattern formation [23, 68–72].

It is still not known whether a Voronoi-like relationship between hydathodes and major veins offers any functional benefits compared to alternative network patterns. Computational modeling may help decipher the trade-offs that occur between different patterns in different environments in terms of hydraulic balance [11, 35, 73, 74], structural support [75], and network robustness [19].

Research on pattern formation has for a long time been dominated by Turing’s reaction-diffusion model [7, 76–78] and Wolpert’s positional information model [79–81], but in plants, a different mechanism, based on regulated auxin transport, as opposed to passive diffusion, is responsible for vein formation [82–85]. Although more mathematical work is needed to analyze the breadth of patterns this mechanism can generate, our model of vascular pattern formation in *Pilea* adds to the already rich tapestry of auxin-driven models.

## Materials and Methods

### *Pilea* growth conditions

*Pilea* plants were purchased from Hicks Nursery, Garden City, NY. Plants were propagated by the provider (Harster Greenhouse, CA) by selecting and raising the strongest runners in the generation every 6 months, and were purchased at relatively the same age. Upon arrival, plants were grown in 6-inch pots in an indoor environment with low indirect light. Plants grown in this indoor condition were used in the initial Voronoi I-I-III tests (*n =* 34 leaves from 6 plants), and were used as the control in the stress experiments. Plants used for stress treatments stayed in the indoor condition for at least 1 week before the treatments started.

For the stress treatments, we exposed 6 *Pilea* plants in each of the three conditions, namely shade (22*^◦^*C, R:FR = 0.4, PPFD = 45), high-light intensity (22*^◦^*C, PPFD = 55– 65), and high-temperature (30*^◦^*C, PPFD = 55), for 5 weeks. Plants in stress conditions were grown on a long-day cycle with 16hr light (7 AM to 11 PM) and 8hr dark in growth chambers. Shade and control plants were irrigated with 250 ml water once per week, except high-light and high-temperature treated plants were watered twice every week to keep the soil at the same approximate water level (See Table S6 for treatment condition details). Plants were then shifted to non-stress indoor condition for 1 week, and the emerging leaves that had developed during the stress treatment were collected for analysis.

### Leaf clearing, staining, and imaging

Fresh leaves were phenotyped with a stereo microscope (Zeiss Stemi 508). Each individual leaf was mounted on a customized transparent acrylic platform and laid flat by a covering acetate sheet. For each leaf, multiple, high-resolution micrographs were taken (camera: Zeiss, Axiocam 208, Color) with the abaxial side up and planarly stitched together with software Panorama Stitcher (v 1.11.1).

To image the vein architecture in cleared leaves, we followed the clearing and staining protocol for studies of leaf architecture [86]. Briefly, fresh *Pilea* leaves were first immersed in Histoclear for 1 hr to remove the cuticle, and then immersed in 5% NaOH solution at 37*^◦^*C for 48 hrs in Petri dishes, with one change of solution at 24 hrs. The leaves were then bleached with sodium hypochlorite (3%) until transparent (2-4 hrs, depending on leaf size). Leaves were then rinsed in distilled water, then dehydrated with 30%, 50%, 70%, 90%, and 100% ethanol for 30 minutes each at room temperature. Cleared leaves were then placed in 0.5% safranin in 100% ethanol for staining overnight. Destaining was done by rinsing the leaf specimen in 100% ethanol and acidifying it with 3–6 drops of 6M hydrochloric acid until the mesophyll tissues appeared pale white. Leaves were rehydrated with 90%, 70%, 50%, 30%, and 10% ethanol for 30 min each at room temperature. Finally, the leaves were mounted in a 10% glycerol medium and scanned with a flat-bed scanner (Epson Perfection V600).

### Tracing and graph conversion of major veins and hydathodes

Tracing was performed on the stitched light microscopy images using a digitizing tablet (Wacom Cintiq Pro 16) and commercial graphic software (SketchbookPro 8.8.5). We formulated tracing rules for the major veins based on *Pilea*’s *brochidodromous* vein architecture [37], which means that the major veins form loops with locally consistent widths and do not end in the parenchyma cells (see Supplementary Text Section 2 for the venation characterization).

#### Major vein tracing rules

We traced veins segment by segment. A segment is a relatively straight piece of vein connected by other veins (major or minor) on both ends. Major veins were segments that have consistent widths and are at least 1.5 times the width of the minor veins that they directly connect to. We also required all major veins to end at a junction with other major veins, to ensure all major veins are part of a closed loop.

#### Hydathode tracing rules

Two observations were used to distinguish hydathodes from ground tissue. First, hydathodes are densely packed with cells, which makes them appear less transparent than ground tissues. Second, in mature leaves, hydathodes are anatomically located at minor vein junctions and can be clearly seen as having star-shaped connections to minor veins (SI Fig. 1). Based on the two observations, we traced out the “shadowy dots” that connect to *≥* 4 minor veins as hydathodes.

Tracing of veins and hydathodes was performed by a single researcher and then reviewed and subsequently edited by two other researchers until consensus was reached (see Fig. S1 for examples of graph extractions superimposed with the microscopy images). Traced images were stored as PSD files and layers were output to vector formats. We used NEFI [87], an OpenCV-based network extraction pipeline, to convert vein tracing to an undirected planar graph, where edges (vein segments) and nodes (junctions and connections of vein segments) were defined from the skeleton of the tracing pattern. Hydathode location extraction was performed with in-house code by finding the center of the traced points (Python 3.8). The extracted vein graph (edges and nodes for vein segments) and hydathodes (a node set of hydathode positions) were then stored together as an undirected graph object in NetworkX (2.7.1, a Python library, for studying graphs and networks) under the same coordinate system, to be quantitatively analyzed for the statistical tests.

### Bayesian generalized linear mixed-effects models

We collected multiple leaves from each *Pilea* plant, which introduced a possible dependency for leaves from the same plants. To control for the random effects from plants when calculating performance differences between hydathodes and reference point sets for our statistical tests, we used linear mixed-effect regression models. For discrete hydathode counts data, we used Poisson mixed-effect regression models.

All mixed-effects models were fitted in a Bayesian hierarchical framework using the Bayesian Model-Building Interface (Bambi 0.9.1 [88]). Bambi generates weakly informative priors for all model variables and uses a Hamiltonian Monte Carlo algorithm to draw from the posterior distributions. For each model between a treatment group (in our case, a different point set, or a stress condition) and the control, we used 5 independent chains of 2000 draws, and we recorded the fraction of total draws that lie between 50% to 100% (the probability of direction, *p_d_*). The variable *p_d_* is an index of effect that represents the certainty with which an effect goes in a particular direction, either positive or negative [89]. We calculated (1 *− p_d_*) as the *p*-value to test statistical significance.

### Detailed Methods for the Voronoi I, II, and III tests

#### Voronoi I: Quantifying local deviation

Proof for the perpendicular bisection corollary of Voronoi centers and edges can be found in Green and Sibson (1978) [90]. When two vein polygons share two or more edges instead of one, we picked a random edge among all edges to be tested.

#### Voronoi II: Jaccard similarity

The Jaccard index measures the proportional area overlap between corresponding polygons of two tessellations, and it offers a unitless and interpretable measure of the alignment between the two tessellations. We computed the J-index between the Voronoi polygons generated using hydathodes as centers and the actual *Pilea* major vein polygons. For this test, we used the full set of hydathodes on the leaf as centers, including both single-polygon hydathodes and multi-polygon hydathodes (hydathodes in polygons that enclose more than one hydathode). We also computed the J-indices between the Voronoi polygons generated using the reference point sets as centers and the actual *Pilea* major vein polygons. For these tests, we generated the reference points in only single-hydathode polygons. For multi-hydathode polygons, we fixed the reference points to the locations of the actual hydathodes. This was done for two reasons: (a) so that the the number of centers (and hence, the number of polygons) was the same across all comparisons; and (b) so that the influence of multi-hydathode polygons towards the J-index was equal for all comparisons.

##### White-noise distorted Voronoi tesellations

We used the J-index between the white-noise distorted Voronoi diagrams and the exact Voronoi diagrams to estimate the magnitude of mismatch between *Pilea* major veins and the exact Voronoi diagram. Voronoi diagrams were computed with the hydathode sets of a leaf sample as the Voronoi centers. Then for each Voronoi vertex, we added a vector of fixed length pointing to a random direction to distort the perfect Voronoi diagram and create a “noisy” version of the exact polygons. We explored noise levels ranging from 1% to 20% the length of the mean polygon axes added to the vertices. For each distorted Voronoi pattern, we applied the Voronoi II test with the exact Voronoi diagram and recorded their J-indices (Fig. S2A, B).

##### Mean-split *k-d* tree algorithm

To create an alternative tessellation to test polygon overlap, we modified the *k-d* tree algorithm [91], which is a space-partitioning data structure, to create a spatial subdivision that: 1) recursively forms bisections with point sets, and 2) is produced hierarchically. In 2D, given a set of points as input, the algorithm recursively finds a partition for the point set with one dimension fixed, and the partition of the other dimension is calculated to be the averaged coordinates of all points in the split zone (Fig. S2C). For *Pilea*, we started with splitting all hydathodes in the direction parallel to the main vein rib. In each step, we iteratively used either the parallel or perpendicular direction to subdivide the point sets, until all partitions contain only one hydathode (Fig. S2D).

#### Voronoi III: Center prediction

We used the corollary of Voronoi property to solve for the best-predicted center coordinates given the polygon edges [44].

Given two adjacent polygons, *P_l_* and *P_m_* with a common edge *E_i_*, the predicted Voronoi centers of *P_l_* and *P_m_* are at (*x̂_l_*, *ŷ_l_*) and (*x̂_m_*, *ŷ_m_*). The two end vertices of *E_i_* are (*u_p_*, *v_p_*) and (*u_q_*, *v_q_*). Let *E_i_* be a segment of the line defined by *y = s_i_ x + b_i_*, where

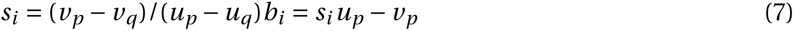

To form a linear system, we will express the Voronoi centers with the geometric relationship in the corollary. The condition on the slope gives:

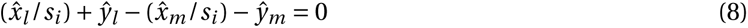

### The condition of distance equality gives

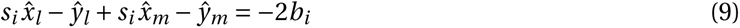

Let (*x_l_*, *y_l_*) and (*x_m_*, *y_m_*) be estimates of Voronoi centers in Eq. (2) and (3). For a tessellation with *k* interior edges and *n* polygons, Eqs. (2) and (3) for all edges together give a system of 2*k* equations with 2*n* unknowns. In ideal Voronoi diagrams, the average number of edges per Voronoi polygon is about 6 [28], therefore 2*k <* 2*n*. Because this test assumes each polygon has exactly one center, we removed from analysis all *Pilea* polygons that contained more than one hydathode. Furthermore, considering a dual graph where nodes are polygons and edges exist between adjacent polygons, we only applied this test to the set of polygons in the largest connected component of this graph. This was done to ensure that there were enough constraints in the system of equations such that the system was not under-determined. Specifically, using the set of polygons in the largest connected component, we found that 33 out of the 34 control samples were over-determined, and hence, we could solve for unique Voronoi center locations by minimizing the *ℓ*_2_ error.

Unlike ideal Voronoi diagrams, the traced *Pilea* vein polygons from tracing sometimes share multiple edges. In such cases, we aimed to find the solution that results in the least distance error with the testing point set (e.g., hydathodes). However, the number of linear systems to select to solve for grows exponentially with the number of polygon pairs that share multiple edges. The heuristic that we adopted for the *Pilea* leaf graphs was to repeatedly pick random edge combinations 100 times the number of polygon pairs, and report the best solution among them. The resulting error is on average 97.8*±*0.7% of the global best solution, computing using brute force on one leaf sample tested, and it performs much better than greedily selecting the best edge (local minimum).

### Sample alignment

To analyze the hydathode distribution from multiple samples, we performed an Ordinary Procrustes style analysis on 32 leaf graphs (8 samples, from each of the 4 environmental conditions). In other words, we aligned all samples in the same orientation and scale by performing isomorphic transformations. The 8 samples we used from each condition were 4 leaves from 2 plants, rather than a random sampling from all leaves, to reduce the variation introduced to the alignment. For every sample, we select the petiole node (*P*) and the furthest intersection node (*Q*) where the primary veins meet as two landmark points. By rotating and scaling the line segment *PQ*s of all samples to perfect alignment, we reach reasonably good superimposition of the leaf graphs, and the hydathode distribution shows a clear spatial pattern, with higher density between the primary and secondary veins that originate from the petiole, and between major secondary veins and the leaf boundary (Fig. 3C).

### Computational modeling

All models were written in C++ extended with constructs of the L+C plant modeling language [92], and executed using the lpfg simulator incorporated into the Virtual Laboratory (vlab) v.5.0 modeling environment (http://algorithmicbotany.org or https://github.com/AlgorithmicBotany). Simulations were performed on a MacBook Pro computer under macOS High Sierra. Videos were assembled from the frames output by lpfg using ffmpeg (https://www.ffmpeg.org).

## Acknowledgments

We thank Kyle Schlecht and Tim Mulligan for plant care and maintenance; Tara Skopelitis, Dr. Thu Tran, and Dr. Fay-wei Li for sharing Marchantia tissue samples; Iacopo Gentile for help designing leaf clearing protocols; Dr. Jia He and Dr. Erika Wee for help with microscopy and imaging; Brendan Brozen for help with tracing; and Dr. Taehoo Ha and Dr. Carlos Martí-Gómez for suggestions on statistical testing. XZ extends her appreciation to thesis committee members, Dr. David McCandlish and Dr. Justin B. Kinney for their support and constructive feedback.

## Author contributions

Conceptualization: XZ, EB, PP, SN

Experiments: XZ, MV, UVP, DJ

Statistical tests & data analysis: XZ, SN

Computational model: PP, XZ

Writing – original draft: XZ, PP, SN

Writing – review & editing: XZ, EB, MV, UVP, DJ, PP, SN

## Data and materials availability

Pre-processed graphs for running statistical analyses are available on Github (https://github.com/cici-xingyu-zheng/VoronoiVein); raw data will be made available by request from the corresponding authors. Code for Voronoi tests can be found on Github (https://github.com/cici-xingyu-zheng/VoronoiVein). All models are available by request from the corresponding authors and will be posted online upon publication.

